# Quantitative Resolving Cell Fate in the Early Embryogenesis of *Caenorhabditis elegans*

**DOI:** 10.1101/2024.10.25.620330

**Authors:** Ruiqi Xiong, Yang Su, Mengchao Yao, Zefei Liu, Jie Lu, Yong-Cong Chen, Ping Ao

**Affiliations:** Shanghai Center for Quantitative Life Sciences & Department of Physics, Shanghai University, Shanghai 200444, China; College of Mechanical Engineering, Beijing Institute of Technology at Zhuhai Campus, Zhuhai 519088, China; College of Biomedical Engineering, Sichuan University, Chengdu 610065, China

**Keywords:** *Caenorhabditis elegans*, embryonic development, endogenous network, landscape, stochastic dynamics

## Abstract

The nematode *Caenorhabditis elegans* exhibits an invariant cell lineage during its development where the gene-molecular network that regulates the development is crucial for the biological process. While there are many molecular cell atlases describing the phenomena and key molecules involved in cell transformation, the underlying mechanisms from a systems biology perspective have received less attention. Based on an endogenous molecular-cellular theory that relates the molecular mechanisms to biological phenotypes, we constructed a model of core endogenous network to describe the early stages of embryonic development of the nematode. Different cell types and intermediate cell states during development from zygotes to founder cells correspond to the steady states of the network as a nonlinear stochastic dynamical system. Connections between steady states form a topological landscape that encompasses known developmental lineage trajectories. By regulating the expression of agents in the network, we quantitatively simulated the effects of the Wnt and Notch signaling pathway on cell fate transitions and predicted the possible trajectories of transdifferentiation of the AB cell across the lineage. The success of the current study may help advance our understanding of the fundamental principles of developmental biology and cell fate determination, offering an effective tool for quantitative analysis of cellular processes.

## Introduction

The development of an organism usually starts with a single cell, where the zygote undergoes numerous divisions to produce millions of cells that differentiate and organize into tissue and organs within the organism. Despite sharing the same genome, these cells differentiate to perform complex and diverse physiological functions. Understanding the genetic regulation of cell fate during embryogenesis is crucial to uncover the general principles of development(Wolpert et al. 2019; Wang et al. 2020). It is believed that some universal developmental principles apply across all animals, making the study of a few model organisms invaluable for elucidating similar processes on a broad range of species. Among them *Caenorhabditis elegans* (*C. elegans*) presents as an important model organism in the developmental biology. It is the only multicellular organism that every cell has been characterized, and it has a persistent cell lineage(Sulston et al. 1983; Kimble and Nüsslein-Volhard 2022) which contains the full set of basic cell types found in higher organisms.

With the completion of the human genome project and the rapid development of high-throughput omics analysis technologies such as single-cell RNA sequencing (scRNA-seq) and cell lineage tracking, the current focus has been shifted focus on to molecular cell maps(Quake 2022). There are numerous studies utilizing scRNA-seq to explore the transcriptome of *C. elegans*. The single-cell transcriptome of the L2 larval stage of *C. elegans* has been well characterized(Cao et al. 2017), along with the construction of cell maps at various stages of early embryonic development(Tintori et al. 2016; Packer et al. 2019; Cole et al. 2024) and the creation of spatiotemporal atlas(Cao 2020; Ma et al. 2021). While classifying cell types based on criteria such as morphology, physiological function, and molecular structure generates vast amounts of data and information(Regev et al. 2017), recording these characteristics for every cell in complex multicellular organisms is impractical. Yet a map as detailed as the reality can be overwhelming and not necessarily helpful(Ao 2008). Therefore it is rather essential to identify some universal approach that can simplify and unify our understanding of these complex biological systems.

During the development phase, cells reach their final differentiated types via distinct fate decisions. Exploring the changes in cell states in the process helps understand the relationship between different cell types. Waddington’s epigenetic landscape conceptualizes developing cells as balls rolling down a hillside, where hilltops represent progenitor cells, and valleys correspond to distinct cell types with varying differentiation potentials(Waddington 1957). The shape of these hillsides is determined by gene regulatory networks(Waddington 1957). For the development process of *C. elegans*, Murray et al. summarized the key regulators of different lineage fates(Murray et al. 2012; Liu and Murray 2023), while Maduro et al. established a gene regulatory network governing cell fate specification in the endomesoderm during the embryonic development(Maduro 2009; Maduro 2010). Du et al. developed a multiscale model that integrates regulatory networks with cell lineage information(Du et al. 2014; Du et al. 2015). Nevertheless, these researches are primarily descriptive, which have yet adequately elucidated the underlying regulatory mechanisms that drive the cell fate decisions. As Sydney Brenner, the Nobel laureate who introduced *C. elegans* to the study of developmental biology, explained in his Nobel Lecture, “We need to turn data into knowledge and we need a framework to do it”(Brenner 2003).

The cell fates of different *C. elegans* lineages are determined at a rather early stage of embryonic development: The zygote undergoes the first asymmetric division to form the anterior AB cell and the posterior P1 cell. The AB cell then divides into ABa and ABp cells, which give rise to hypodermis, neurons, muscle, and other specialized cells. The P1 cell divides to form the EMS and P2 cells. The EMS cell leads to the endomesoderm of *C. elegans* and further divides into MS cells. The latter produce muscle, glands, coelomocytes, and E cells which form the intestine. The P2 cell undergoes two additional divisions, forming C and D cells, which produce various tissues along with the P4 cell that creates the germ cells(Wolpert et al. 2019). Although *C. elegans* has a consistent cell lineage, where cell fate at each stage of development is well-documented, this does not imply that cell differentiation always follows the same trajectory(Rothman and Jarriault 2019) due to the complex interplay among genes, proteins, and cells that drives the embryonic development. Thus the cell fate decisions during the early embryonic development of *C. elegans* are ideal for studying how the interactions of these factors set the development.

In this study, we have constructed the core endogenous network of early embryonic development of *C. elegans* in which the interactions between proteins and genes are modeled as a high-dimensional nonlinear stochastic dynamical system. Computational analysis recovers the steady states that correspond to the expression profiles of the five founder cells (AB, MS, E, C, D), the stem cells (P1–P4), and several daughter cells (ABa, ABp, Z2/Z3), along with a series of transition states representing intermediate cell states. By tracing the evolutionary trajectories from transition states to steady states, the topological landscape of the early embryonic development of *C. elegans* can be mapped, which encompasses the normal developmental pathways as well as raises transition pathways between cells that are not documented. These results are independent of high-throughput data, allowing for the prediction of intermediate cell states and transdifferentiation pathways yet observed experimentally. Furthermore, by regulating the expression of specific agents, cell transitions between different states can be induced, offering a foundation in silico to extend the framework, potentially offering a quantitative approach for studying, for instance, animal development in other species.

## Materials and methods

### 1. Endogenous network construction

The endogenous network modeling proposed by Ao, Hood and *et al*. presents a alternative approach to characterize complex molecular regulatory mechanisms in biological systems from a systems biology perspective, facilitating quantitative analyses through stochastic dynamics(Ao et al. 2008). The theory posits that an endogenous molecular-cellular network, shaped by billion years of biological evolution, can be quantitatively represented by the expression or activity levels of key endogenous agents, forming a high-dimensional stochastic dynamical system. Nonlinear interplay and dynamical interactions between the network agents lead to multiple local stable states, each with distinct biological functions. The network dynamics autonomously achieves and maintains these steady states over extended periods. The inherent stochasticity in biochemical reactions channels transitions across the landscape barriers between steady states, corresponding to cell state changes during differentiation. Identifying key agents within the endogenous network, detailing their interactions, and elucidating their global dynamical properties can systematically uncover the mechanisms underlying the normal and abnormal tissue functions(Yuan et al. 2017).

To construct an endogenous network for a specific study, one would first select modules based on the biological phenomena and functions to be described. These functional modules are inseparable but relatively autonomous components of a biological system, which can either perform modular functions independently or jointly execute cell fate decisions through complex crosstalk(Hartwell et al. 1999). In this study, we would focus on the early embryonic development stage, specifically the initial cell divisions from fertilization to the generation of the founder cells(Wood 1988). Based on the functional roles of cells at this stage, we selected four key modules, cell polarity, cell cycle, signaling pathways, and lineage specification with the core agents for each module and the interactions between them. An agent (i.e. node) on the network, in the current context, can represent typically a cluster of relevant genes / molecules and/or a signaling pathway. These genes or proteins regulate each other directly or indirectly through transcription, translation, or signaling, hence we take a coarse-grained approach to abstract these regulations into “activation” or “inhibition” interactions.

The polarity during asymmetric cell division is primarily controlled by the PAR polarity proteins(Bowerman et al. 1997). In what follows PAR-6 represents the anterior protein complex PAR-6/PAR-3/PKC-3, while LGL-1 represents the posterior complex LGL-1/PAR-1/PAR-2(Hoege and Hyman 2013; Rose and Gönczy 2014) and PLK-1 stands for the complex PLK-1/MEX-3/MEX-5, which links polarity protein to the asymmetric distribution of agents within the cytoplasm(Budirahardja and Gönczy 2008). The daughter cells of AB divided more rapidly than those of P1. Cdc-25,1 and CDK-1 are chosen to represent the promoters of the cell cycle, while WEE-1.1 represents the inhibitors (Van Den Heuvel 2005; Tavernier et al. 2015; Kipreos and Van Den Heuvel 2019). During the division of AB and EMS to generate daughter cells, two key intercellular signal transductions are involved. GLP-1, a receptor, represents the Notch pathway, and the TCF transcription factor POP-1 represents the Wnt signaling pathway (Mello et al. 1994; Phillips et al. 2007). Key factors that determine cell differentiation include maternal transcription factors like SKN-1 for mesoderm and endoderm specification(Bowerman et al. 1992; Bowerman et al. 1993; Gilbert and Barresi 2016), PIE-1 for germline cell fate(Mello et al. 1996; Gilbert and Barresi 2016), and PAL-1 for the somatic descendants of the P2 blastomere(Hunter and Kenyon 1996; Edgar et al. 2001; Gilbert and Barresi 2016). Additionally, lineage-specific factors include TBX-35 and TBX-37 for mesoderm(Good et al. 2004; Broitman-Maduro et al. 2006), PHA-4 for the pharynx(Kalb et al. 1998; Kiefer et al. 2007), HLH-1 for muscle(Fukushige and Krause 2005; Fukushige et al. 2006), END-3 for intestinal endoderm(Zhu et al. 1997; Ewe et al. 2022), and LIN-26 for ectoderm(Labouesse et al. 1996). The above agents are listed in Table S1 in Supplementary Materials along with their associated modules. The mutual interactions are further illustrated in Figure 1a.

**Figure 1.**
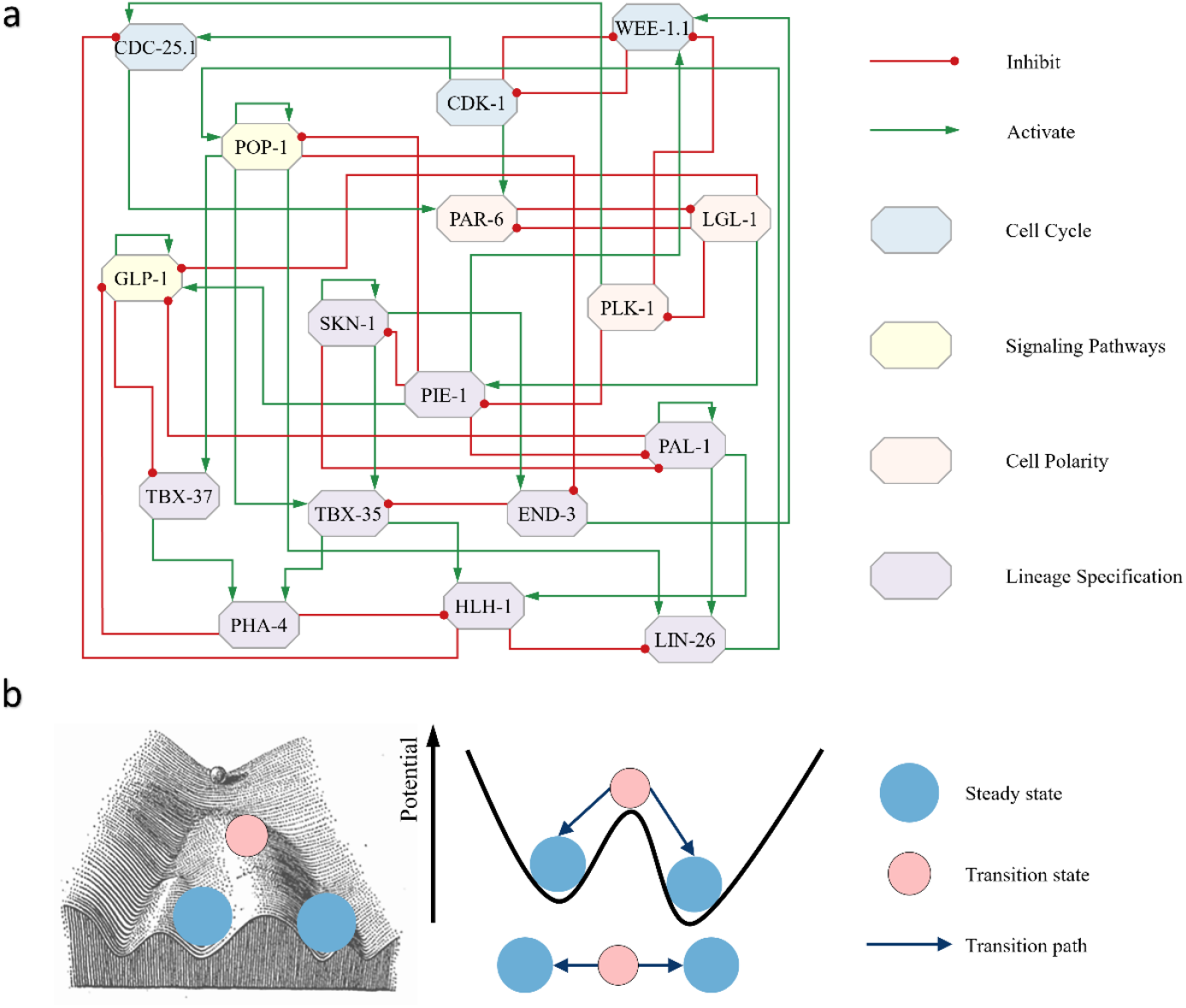
a) Schematic representation of the core endogenous network of early embryonic development of C. elegans. The network has 17 nodes and 43 edges. Blue nodes are selected from the cell cycle module, while yellow colors are for signaling pathways, pink ones are agents from the cell polarity module, and finally purple nodes are selected for the lineage specification. b) Potential illustration of Waddington’s epigenetic landscape. The noise-driven state transition is bidirectional and represented by the double arrows in the figure.

### 2. Nonequilibrium stochastic dynamics

The dynamics of *N* agents within the endogenous network, with their level of expressions or activities denoted as **x** = (*x*_1_, *x*_2_, …, *x*_*N*_)^*T*^, can be represented by a set of stochastic differential equations (SDEs):

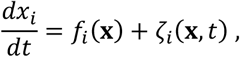

where the deterministic part of the “force” *f*_*i*_(**x**), a generically nonlinear function is the rate of change for the expression level *x*_*i*_ of the *i*^th^ agent, *ζ*_*i*_ represents multiplicative Gaussian white noise with ⟨*ζ*_*i*_(**x**, *t)*⟩ = 0, ⟨*ζ*_*i*_(**x**, *t)ζ*_*j*_(**x**, *t*′)⟩ = 2ε*D*_*ij*_(**x***)δ*(*t* − *t*′). Here ε indicates the noise intensity, ⟨⋯ ⟩ stands for the average over the fluctuations, *D*(**x**) denotes the diffusion matrix, and finally *δ*(*t)* is the Dirac delta function. Such a general form of SDE can be further decomposed into the following form(Ao 2005),

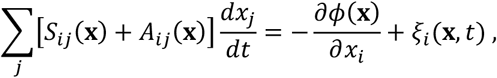

where the friction matrix *S*(**x**) corresponds to dissipation, the transverse matrix *A*(**x**) corresponds to oscillations, the scalar function *ϕ*(**x**) represents a generalized potential, and *ξ*_*i*_ is “normalized” zero-mean white noise whose covariance satisfies ⟨*ξ*_*i*_(*x, t)ξ*_*j*_(*x, t*′)⟩ = 2ε*S*_*ij*_(**x***)δ*(*t* − *t*′). By substituting the first set of equations into the decomposed form and equating the deterministic and stochastic parts separately, we get a relationship between the friction and the transverse matrices to the diffusion matrix and the force. The potential function can be nominally calculated via *ϕ*(**x**) = − ∫[*S*(**x**) + *A*(**x**)]*f*(**x***)d***x**. Near a local minimum, the expression level has a Boltzmann-like distribution *ρ*(**x**) ∝ exp(−*ϕ*(**x**)/ε). It can be further extended to steady-state distribution without detailed balance. The existence of such potential function (or the “energy” in the current context) sets a solid foundation for the subsequent landscape analysis.

The fixed points of the steady-state distribution calculated using this method coincide with the fixed points of the ordinary differential equations (ODEs) in the absence of the noise, allowing quantitative analysis on the endogenous network with dimensions ranging from tens to hundreds. Taking now the deterministic part of the equation *f*_*i*_(**x**), the effect of activation and inhibition of other agents on the rate of the *i*^th^ agent can be simulated using a Hill function of the form,

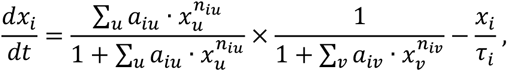

where the exponent *n* is the Hill coefficient, *a* is the inverse of the apparent dissociation constant, and the summation 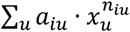 and 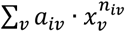 are over the the activation and inhibition agents respectively. In the current context, *n* describes the cooperative interactions among agents in the dynamics, and τ denotes some generic degradation constant. With the difficulty of obtaining precise values of these parameters across several orders of magnitude, we focus on the relative degrees of activation and expression between agents, i.e., the relative activation or expression levels for each agent is normalized to 0-1, with the associated degradation constant τ set to 1. To confine the threshold of the Sigmoid form of the Hill function to around 0.5 thus capturing the key characteristics of the dynamics, we set the constraint *a* = 2^*n*^, with *n*=4, *a*=16 in the simulations in this work.

In the extreme limit where *n* approaches infinity, the differential equation transitions from a Sigmoid form to a step function, converted to a discrete set of Boolean operations. For instance, the Boolean expression for the regulatory relationship of the agent PAR-6 in the network is given as PAR-6(t+1) = (CDC-25.1(t) OR CDK-1(t)) AND (NOT LGL-1(t)). Boolean dynamics is a powerful tool for capturing the overall structural information of a network, providing a robust approximation of nonlinear dynamical behavior. This approach has broad applications in the description and analysis of complex biological systems(Bornholdt 2008; Albert and Thakar 2014). Comparing ODE dynamics with Boolean dynamics allows for the testing of both methods’ validity and the identification of saddle points that Boolean dynamics alone cannot compute. It reaffirms the hypothesis that the system’s dynamic structure at the core endogenous network level is largely parameter-insensitive, i.e. determined primarily by the network’s topology.

### 3. Fixed-Points of the ODEs

Two distinct methods were used to identify the fixed points of the nonlinear ODEs.

#### Algorithm 1

Starting with a randomly generated *N*-dimensional initial state **x**_0_, the evolution of the state vector is updated with iteration via *x*_*t*+1_ = *x*_*t*_ + Δ*t × f*(*x*_*t*_). After a large number of operations *N*_*t*_, the network reaches a steady state when (*x*_*t*+1_ − *x*_*t*_)^2^ *<* ε, *f*(*x*_*t*_)^2^ *< δ* with sufficiently small ε and *δ*. In this work, we set Δ*t* = 0.1, *N*_*t*_ = 800, ε = 10^−8^, *δ* = 10^−8^. Repeat the process until all steady states in the phase space are obtained.

#### Algorithm 2

Use the Newton’s iteration method to solve the nonlinear equations *f*_*i*_(**x**) = 0, *i* = 1,2 ⋯,N, which identifies both stable and unstable fixed points in the system. Starting again with random initial vector **x**_0_, the method finds *x*^∗^ that satisfies *f*_*i*_(*x*^∗^) = 0. This process can be implemented using MATLAB’s fsolve function. When real parts of the eigenvalues of the Jacobian matrix at *x*^∗^ are all negative, it is recorded as a stable fixed point, i.e., a steady state. Otherwise the solution is unstable fixed point, i.e. a saddle point or a transition state. The number of eigenvalues with positive real part is also recorded, and a transition state with more than such eigenvalues is referred to as a hyper-transition state. Repeating the process for example, 10^6^ times or more, to exhaust the search of fixed points in the phase space.

Having identified the system’s stable and transition states, fluctuations are applied to obtain the paths from each transition state to the steady states nearby. Namely, add a small random vector Δ**x** to *x*^∗^ as a local perturbation, let the system iterate under *x*_*t*+1_ = *x*_*t*_ + Δ*t × f*(*x*_*t*_), then record the evolution path from the perturbed transition state to the final steady state. Again the dynamics is converged when (*x*_*t*+1_ − *x*_*t*_)^2^ *<* ε, *f*(*x*_*t*_)^2^ *< δ*. Then, we can obtain the steady states and transition paths in the landscape determined by the endogenous network, as shown in Figure 1b.

## Results

### 1. Endogenous network simulation of *C. elegans*

To quantitatively explore the mechanisms of cell fate decisions during early embryonic development of *C. elegans*, we constructed a core endogenous network specific to this phase, with the relevant modules and core factors involved and the interactions between them shown in Figure 1a. It is a closed system where each agent is regulated by other agents, forming feedback loops with no pure upstream-downstream relationship. It has the ability to make its own decisions and can evolve over time to stable states with distinct biological phenotype.

As described in the Methods section, the network dynamics can be modeled by a set of nonlinear ODEs for analysis. Carrying out the simulations we obtained 10 steady states and 33 transition states, with their corresponding expression profiles displayed in Figures 2a and 2b, respectively. On a separate count, Boolean dynamics for initial state traversal gives rise to 18 attractors, including 10 point attractors and 8 linear attractors, as illustrated in Figure 2c. The 10 point attractors match the 10 steady-state profiles obtained through ODE analysis. Additionally, the linear attractors correspond to a subset of the transition states, confirming the consistency between these two computational approaches as discussed in the Methods section.

**Figure 2.**
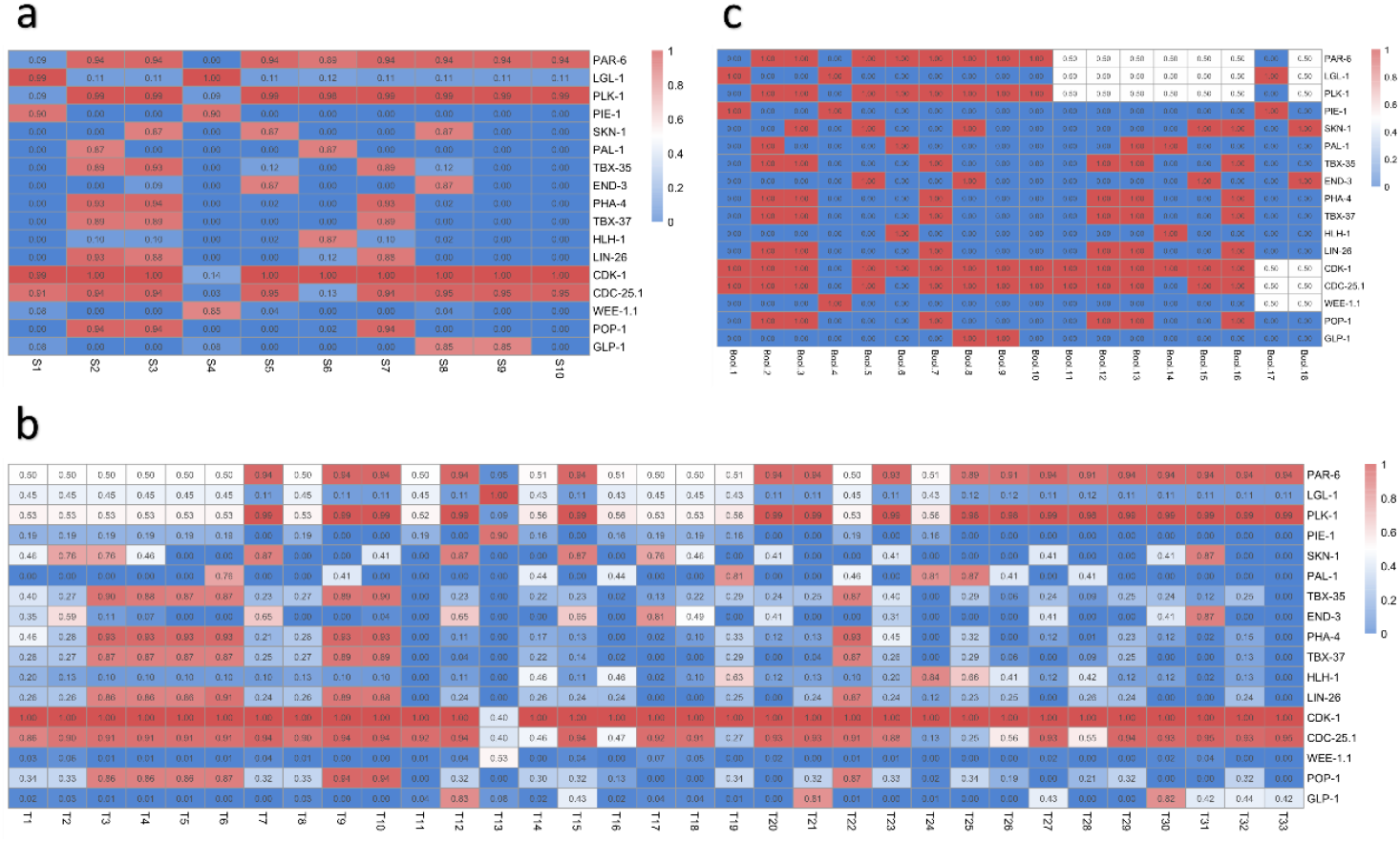
a) The 10 steady states obtained from ODE. b) The 33 transition states identified by ODE. c) The 10 point attractors and 8 linear attractors based on Boolean dynamics. Red colors in the grids mark high expression levels and blue for low expression ones.

### 2. Phenotypes identified with experimental data

The biological characteristics for each steady state was initially determined based on the expression of specific marker factors associated with each cell type. Both S1 and S4 (steady states) exhibited high expression of the posterior factor LGL-1 and the germline factor PIE-1, indicating their correspondence to the P lineage cells. Furthermore, in S1, cell cycle-promoting factors are highly expressed while the levels of inhibitory factors remain low, whereas in S4, this pattern is reversed. Thus, S1 is identified as P1-P4 cells, which continuously divide with stem cell characteristics during early development, while S4 corresponds to Z2/Z3 cell that enters a quiescent state during late embryonic development(Schedl 2013). In S3, S5 and S8, the endomesoderm activator SKN-1 is highly expressed. Additionally, the mesodermal factor TBX-35 is highly expressed in S3. Therefore, it can be determined that S3 corresponds to the MS cell that mainly divides to generate mesoderm. The endodermal factor END-3 is highly expressed in both S3 and S8. Considering the transition pathways between steady states, only S5 can be directly converted from S1, representing the P lineage, through a single transition state. Therefore, S5 is identified as the EMS cell, and S8 as the E cell, which develops into the intestine. PAL-1 is highly expressed in both S2 and S6, suggesting that these states correspond to C and D cells, which generate somatic cells in the P2 daughter cells. Since D cells differentiate exclusively into muscle(Gilbert and Barresi 2016), S6, with its high HLH-1 expression, is identified as the D cell, while S2 is identified as the C cell. In S7, S9, and S10, the anterior factors PAR-6 and PLK-1 are highly expressed, and the maternal transcription factors SKN-1, PIE-1, and PAL-1, which are asymmetrically distributed in P1, are expressed at low levels, indicating that these states correspond to the AB lineage. The endodermal factor TBX-37 is highly expressed in S7, and the Notch ligand GLP-1 is highly expressed in S9, leading to the identification of S7 as ABa, S9 as ABp, and S10 as general AB cells. The results of low-throughput experiments from the literature, summarized in Figure 3a, are consistent with our calculated outcomes.

**Figure 3.**
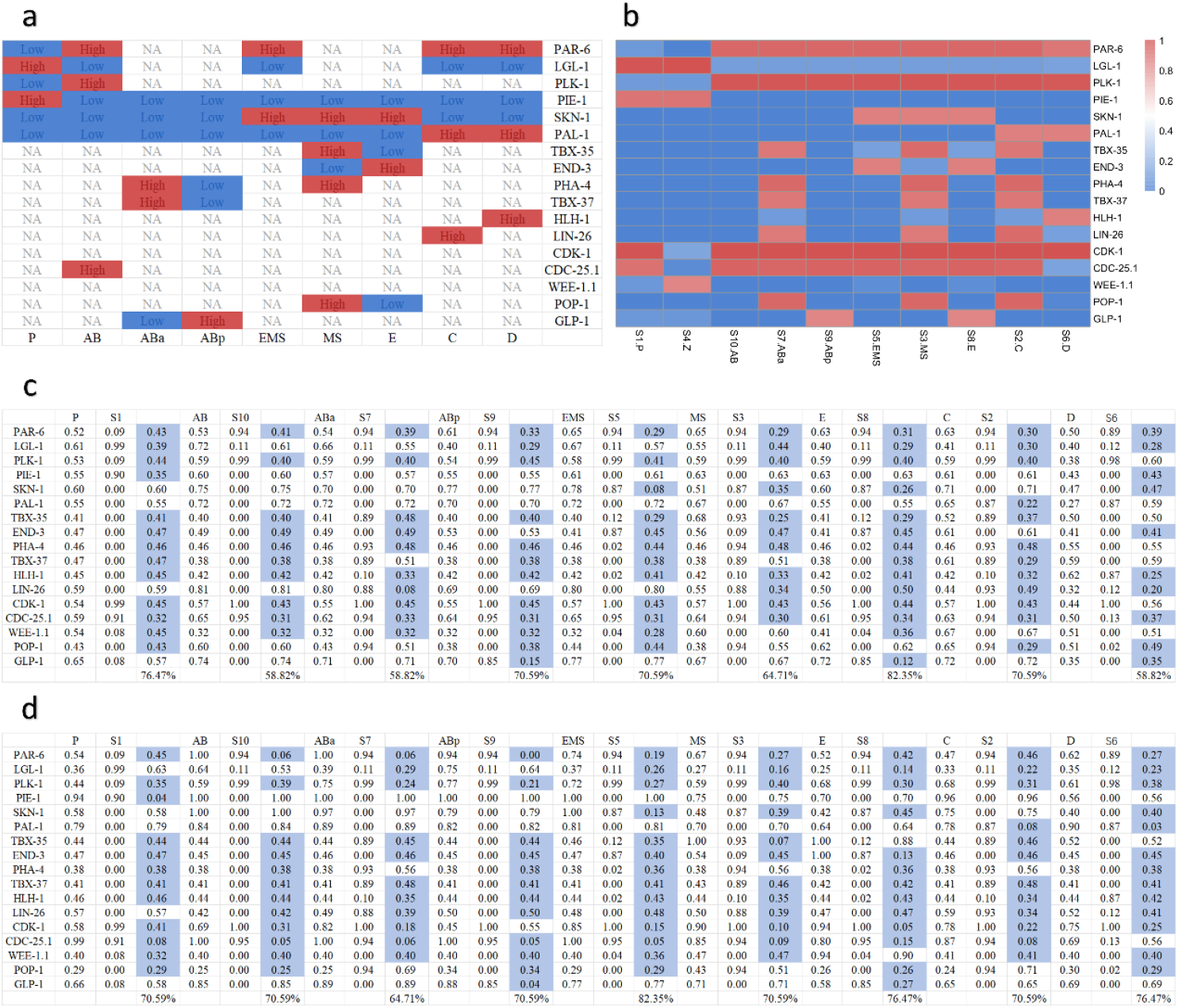
a) Low-throughput data collated from the literature. b) Comparison between steady-state and low-throughput data of the 1-16 cell stage. c) The comparison between steady-state and high-throughput data. The first column is the normalized high-throughput data, the second column is the simulated results, the third column is the difference between the two, and so on. The purple (light-blue) grids are for those with less than 0.5 difference, i.e. the calculated results are consistent with the experimental data, with 58%-82% consistency rates. d) The comparison with the experimental data of the 1-102 cell stage, with 64%-82% consistency rates.

The steady-state expression profiles were further validated by comparison with high-throughput data. Two datasets, GSE77944 and GSE83523, were used, which measure single-cell transcriptome expression in *C. elegans* embryos as they develop from the 1-cell stage of a fertilized egg to the 16-cell and 102-cell stages, respectively(Tintori et al. 2016; Cole et al. 2024). The data are normalized, with 0 representing the lowest expression of the factor and 1 representing the highest expression. A threshold of 0.5 was established, where differences less than 0.5 were considered consistency, and differences equal to or greater than 0.5 were considered inconsistent, and the consistency rates between the calculated steady states and the experimental data were found to range from 58%-82% for the first dataset and 64%-82% for the second dataset, as illustrated in Figure 3c and 3d. It is important to note that the factors with less consistency across both datasets were SKN-1, PIE-1, PAL-1, and GLP-1. These discrepancies likely arise because the network calculations represent the concentration of proteins in their active states, while SKN-1, PIE-1, and PAL-1 are maternal transcription factors whose regulation occurs primarily at the levels of translation and post-translational modification(Baugh et al. 2003; Gilbert and Barresi 2016), leading to differences between mRNA expression levels and actual protein concentrations. GLP-1, a receptor located on the cell membrane(Evans et al. 1994; Priess 2005), has an active concentration that does not necessarily correlate with its mRNA expression levels. In conclusion, the compliance ratio between the calculated steady states and the experimental data fell consistently within a reasonable range, strongly supporting the correspondence of these steady states to the cell types observed during early embryonic development of *C. elegans*.

### 3. Topology of the steady-states of landscape

To characterize the process of cell fate determination, we further applied local perturbations to the network to investigate the interconnections between different steady states and transition pathways, as detailed in the Methods section. Putting together the results of perturbations across all transition states and comparing the post-perturbation steady states with those obtained earlier, we obtained a landscape topology of the endogenous network constructed, as depicted in Figure 4a. Here the order of the hyper-transition states corresponds to the number of eigenvalues of the Jacobian matrix having non-negative real part. The landscape allows us to interpret the role of regulation of transcription factors during the development. It facilitates the transition of cells between different steady states by overcoming potential barriers at the intermediate states, which are shaped by the transcription factors.

**Figure 4.**
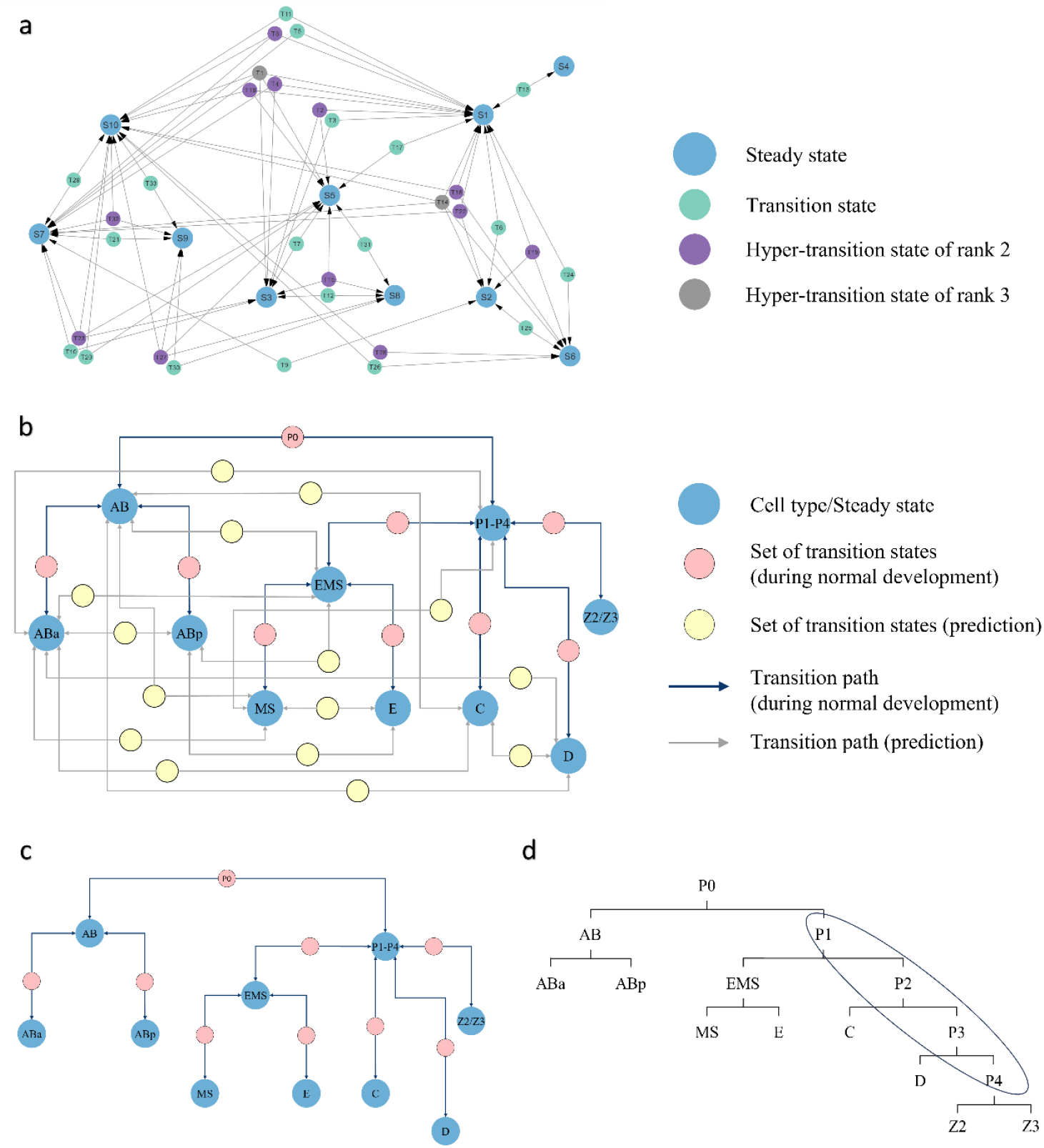
a) Transition paths between steady states on the endogenous network. b) Landscape of early embryonic development of C. elegans constructed from the paths. c) Part of the computed landscape that matches the observed developmental trajectory. d) Lineage of early embryonic development of C. elegans. The circled portion is the P1-P4 cell types that is grouped into the same steady state in the modeling.

We classified the transition states into different sets in accordance with the connectivity between the steady states and summarized the transition paths between steady states that emerge through these sets of transition states, as detailed in Table 1. Consequently, we were able to construct the landscape of early embryonic development of *C. elegans*. This outcome represents a quantitative realization of Waddington’s epigenetic landscape, encompassing 10 experimentally observed cell types: AB, ABa, ABp, EMS, MS, E, C, D, P, and Z, as well as possible switching pathways between these cell types connected by transition states. The constructed landscape reveals 24 developmental pathways during early embryonic development, 9 of which align with experimentally observed phenomena and correspond to the trajectories of the cell state changes during the first few cleavages of early embryonic development of *C. elegans*, as shown in Figure 4c. Note that a single transition state can connect multiple steady states, presenting intersections between the pathways.

**Table 1.**
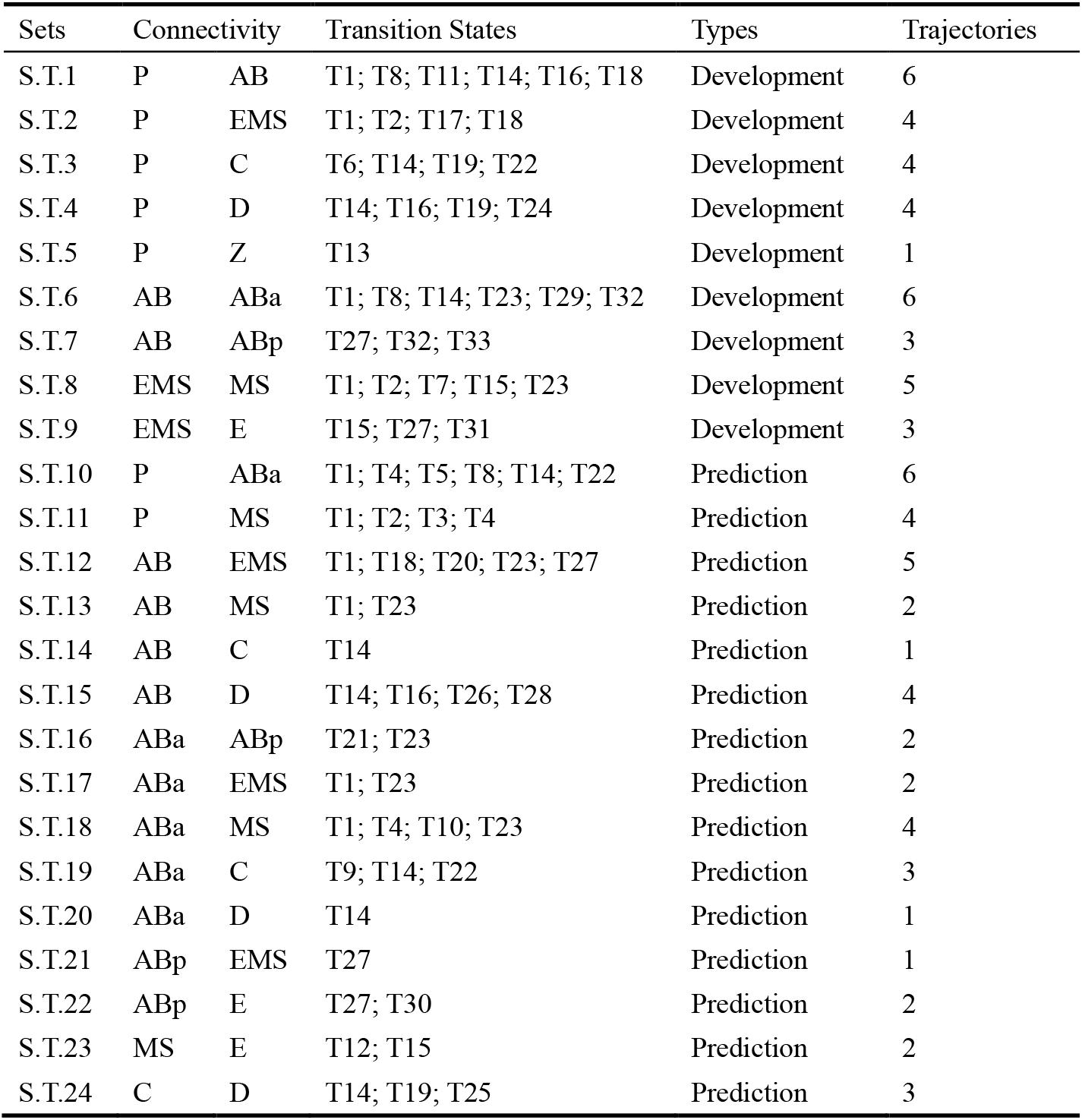
Developmental pathways in the landscape of early embryonic development of C. elegans. There are 24 sets consisting of transition states. Among them 9 matched to developmental processes and the rest are predicted by the simulation. There can be multiple routes in each site, resulting in total 78 different trajectories.

### 4. Simulation of cell fate regulation

Transitions between different cell states can be simulated by modulating the expression levels of specific agents within the network. First, one can emulate the effects of intercellular signaling pathways. At the 4-cell stage, the P2 cell transmits a Notch signal to the AB cell, leading to the activation of the receptor GLP-1 on the membrane surface of the ABp cell, which is the daughter cell of AB in contact with P2. This signaling inhibits the development of the ABp cell’s daughter cells into mesoderm(Priess 2005). In our model, steady state S10 represents the AB cell. To simulate the signaling effect of the P2 cell, we increased the expression of the germline-specific factor PIE-1 in the AB cell’s expression profile. The evolutionary pathway from this initial state proceeds through the transition state T33 en route to the steady state S9, which represents the ABp cell, consistent with the phenomenon during development. Similarly, Wnt signaling from P2 cell to EMS daughter cell promotes the fate of E cell(Owraghi et al. 2010). By increasing the expression of PIE-1 in the steady state S5, representing the EMS cell, the evolutionary pathway progresses through the transition state T31 then into the steady state S8 representing the E cell. These results shown in Figure 5a are consistent with the observed developmental processes, which further validates the core endogenous network model.

**Figure 5.**
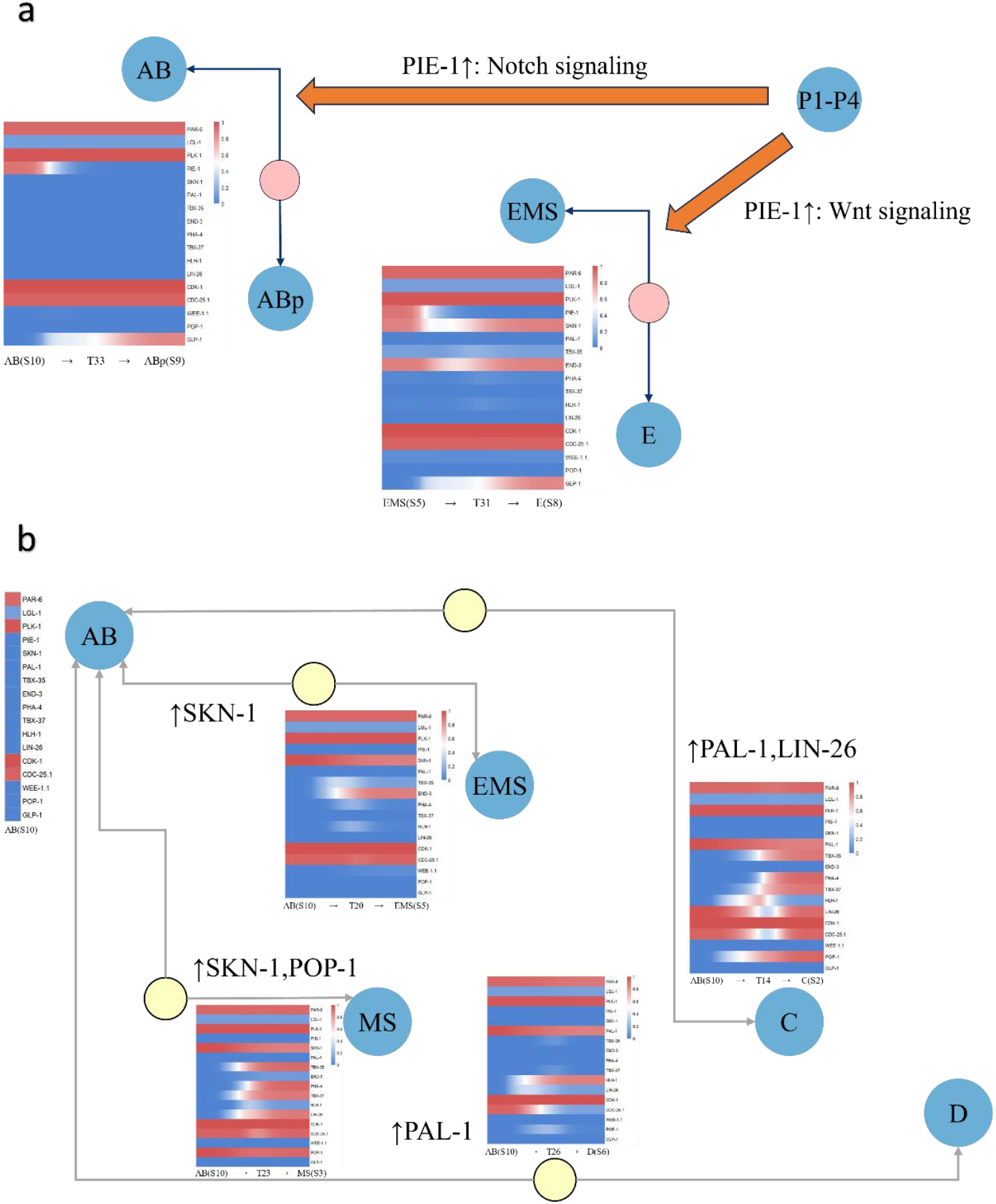
a) The signaling effects on P2 cell, simulated by up-regulating the expression of PIE-1, along the trajectories upon receiving the signal as AB and EMS cells divide into daughter cells. b) Predicted possible regulatory approaches that can induce trans-lineage switching from AB cell to other cell types.

We further explored the transition pathways in the vicinity of AB cells that do not occur during normal development. Overexpression of SKN-1 in steady state S10, representing the AB cell, moves the system to steady state S5, representing the EMS cell, via transition state T20. When both SKN-1 and POP-1 are overexpressed, it goes through the hyper-transition state T23 to the steady state S3 representing the MS cell. Similarly, overexpression of PAL-1 in S10 leads to its conversion via transition state T26 to steady state S6, representing the D cell. When both PAL-1 and LIN-26 are overexpressed, S10 transforms into steady state S2, representing the C cell, via hyper-transition state T14, as shown in Figure 5b. These findings predict potential regulatory strategies for inducing AB cells to switch to alternative lineages. They will further be discussed below.

## Discussion

The early embryonic development of *C. elegans* has been extensively studied, offering a wealth of experimental data that provides an ideal platform for further analysis. Based on extensive search over the literature, we selected 17 core agents, along with their promoting or inhibiting interactions to construct the core endogenous network of early embryonic development of *C. elegans*. The evolutionary dynamics of the network naturally gives rise to distinct cell fates and various intermediate states.

During the initial divisions of the zygote, each asymmetric division generates a progenitor cell, which can produce somatic descendants, and a stem cell (P1-P4 cells) (Schedl 2013; Gilbert and Barresi 2016). In this study, we classify P1-P4 cells to the same steady state S1 in our modeling. From the perspective of network dynamics, P1-P4 cells are self-renewing in the same cellular state, they produce the progenitor cells which differentiate into distinct steady states. The transition pathways linking these steady states reflect the developmental landscape of the cells. In Figure 4, the steady state representing germ cells Z2/Z3 cannot transition into the somatic cell developmental pathway via a single transition state, in consistence with the separation of somatic and germ lineages during embryonic development(Seydoux and Fire 1994).

The pathways within the landscape generated by the endogenous network are bidirectional. For instance, the transition pathways between P cells and somatic cells can represent both the differentiation of P cells into somatic cells through a transition state and a reversal of the latter back to the germline stem cells via the same transition state, suggesting a potential for somatic cells to de-differentiate. Beyond the pathways observed during normal embryonic development, the landscape reveals additional transition pathways, as illustrated by the gray line in Figure 4b. Among them the conversion involving the AB lineage is most extensive, corresponding to the broad distribution of daughter cells of AB lineage across the neuronal systems, pharynx, skin, body wall muscle, and others. These pathways possess significant differentiation potential(Sulston et al. 1983). Moreover, the presence of multiple pathways between these stable states reflects redundancy and robustness in the development. Namely the same cell type can be produced even after variations in regulation or gene mutation(Maduro et al. 2005; Fukushige et al. 2006). It appears that the outcomes of cell fate decisions rely more on the genomic blueprint than on the specific developmental processes.

By quantitatively regulating some target agents, one can model how intercellular signaling impacts cell states, leading to transdifferentiation of the AB cell through non-canonical pathways. Along with the temporal dynamics of core endogenous agents during these transitions, it sets up theoretical foundation for induced regulation of cell states. Such research can be of great importance in developmental biology, for example can study how cells achieve specific fate decisions in specific environments through delicate signaling regulation. It may identify key disease-related factors, offering potential targets for precise interventions and drug therapies(Zhang et al. 2024), thereby advancing the field of precision medicine in a broad perspective.

Based on the core endogenous network, the topological landscape obtained has successfully characterized the early embryonic development of *C. elegans*. Nevertheless, the current focus on modules and factors involved in the first few cleavages limits our exploration for later stages, restricting the scope to the earlier progenitor cell stages of each lineage. With the incorporation of apoptosis and other key modules, works may be extended to the developmental processes of each lineage during the gastrulation stage and beyond. This can cover the dynamic mechanisms responsible for generating diverse cell types, tissues, and organs throughout development.

## Conclusion

To explore the mechanisms underlying the developmental progress from the perspective of systems biology, a model organism, *Caenorhabditis elegans*, was analyzed in the platform of endogenous network for its early embryonic development. By encapsulating genome-level interactions and cell-level phenotypic data within a causal-dynamical framework, we were able to recover all the observed differentiation processes as well as raise further predictions awaiting experimental validation. With the current success, similar studies in more complex organisms may also be carried out. This work demonstrates the high applicability of the theoretical groundwork adopted in this research.

## Data Availability Statement

Single-cell transcriptome data of *C. elegans* cells in early blastomeres up to the 16-cell stage can be accessed in the GEO database under the code GSE77944. The transcriptomes up to 102-cell stage can be accessed in the GEO database under the code GSE83523.

## Acknowledgement

This work was supported in part by National Natural Science Foundation of China (Y.-C.C., Approval No. 12375034).

## Author contributions

Conceptualization, R.X., Y.S., M.Y., Y.-C.C. and P.A.; methodology, Y.-C.C. and P.A.; investigation, R.X., Y.S.and M.Y.; resources, J.L., Y.-C.C and P.A.; data curation, R.X., Y.S., M.Y. and Z.L.; writing—original draft preparation, R.X. and Z.L.; writing — review and editing, R.X., Y.S., M.Y., Z.L., J.L., Y.-C.C. and P.A.; project administration, J.L., Y.-C.C. and P.A.; funding acquisition, Y.-C.C..

## Additional information

Supplementary Materials

## Reference

Albert R, Thakar J. 2014. Boolean modeling: a logic-based dynamic approach for understanding signaling and regulatory networks and for making useful predictions. WIREs Syst Biol Med. 6(5):353–369. doi:10.1002/wsbm.1273.

Ao P. 2005. Laws in Darwinian evolutionary theory. Phys Life Rev. 2(2):117– 156. doi:10.1016/j.plrev.2005.03.002.

Ao P. 2008. Borges Dilemma, Fundamental Laws, and Systems Biology.Bioinforma Biol Insights. 2:201–2. doi:10.1177/117793220800200002.

Ao P, Galas D, Hood L, Zhu X. 2008. Cancer as robust intrinsic state of endogenous molecular-cellular network shaped by evolution. Med Hypotheses. 70(3):678–684. doi:10.1016/j.mehy.2007.03.043.

Baugh LR, Hill AA, Slonim DK, Brown EL, Hunter CP. 2003. Composition and dynamics of the Caenorhabditis elegans early embryonic transcriptome. Development. 130(5):889–900. doi:10.1242/dev.00302.

Bornholdt S. 2008. Boolean network models of cellular regulation: prospects and limitations. J R Soc Interface. 5(Suppl_1). doi:10.1098/rsif.2008.0132.focus. [accessed 2024 Aug 31]. https://royalsocietypublishing.org/doi/10.1098/rsif.2008.0132.focus.

Bowerman B, Draper BW, Mello CC, Priess JR. 1993. The maternal gene skn-1 encodes a protein that is distributed unequally in early C. elegans embryos. Cell. 74(3):443–452. doi:10.1016/0092-8674(93)80046-H.

Bowerman B, Eaton A, Priess JF. 1992. skn-1, a Maternally Expressed Gene Required to Specify the Fate of Ventral Blastomeres in the Early C. elegans Embryo. Cell. 68(6):1061–75. doi:10.1016/0092-8674(92)90078-Q.

Bowerman B, Ingram MK, Hunter CP. 1997. The maternal par genes and the segregation of cell fate specification activities in early Caenorhabditis elegans embryos. Development. 124(19):3815–3826. doi:10.1242/dev.124.19.3815.

Brenner S. 2003. NATURE’S GIFT TO SCIENCE. Biosci Rep.(5–6):225–232. doi:10.1023/B:BIRE.0000019186.48208.f3.

Broitman-Maduro G, Lin KT-H, Hung WWK, Maduro MF. 2006. Specification of the C. elegans MS blastomere by the T-box factor TBX-35. Development. 133(16):3097–3106. doi:10.1242/dev.02475.

Budirahardja Y, Gönczy P. 2008. PLK-1 asymmetry contributes to asynchronous cell division of C. elegans embryos. Development. 135(7):1303–1313. doi:10.1242/dev.019075.

Cao J. 2020. Establishment of a morphological atlas of the Caenorhabditis elegans embryo using deep-learning-based 4D segmentation. Nat Commun. 11(1). doi:10.1038/s41467-020-19863-x.

Cao J, Packer JS, Ramani V, Cusanovich DA, Huynh C, Daza R, Qiu X, Lee C, Furlan SN, Steemers FJ, et al. 2017. Comprehensive single-cell transcriptional profiling of a multicellular organism. Science. 357(6352):661–667. doi:10.1126/science.aam8940.

Cole AG, Hashimshony T, Du Z, Yanai I. 2024. Gene regulatory patterning codes in early cell fate specification of the C. elegans embryo. eLife. 12. doi:10.7554/eLife.87099.2.

Du Z, Santella A, He F, Shah PK, Kamikawa Y, Bao Z. 2015. The Regulatory Landscape of Lineage Differentiation in a Metazoan Embryo. Dev Cell. 34(5):592– 607. doi:10.1016/j.devcel.2015.07.014.

Du Z, Santella A, He F, Tiongson M, Bao Z. 2014. De Novo Inference of Systems-Level Mechanistic Models of Development from Live-Imaging-Based Phenotype Analysis. Cell. 156(1–2):359–372. doi:10.1016/j.cell.2013.11.046.

Edgar LG, Carr S, Wang H, Wood WB. 2001. Zygotic Expression of the caudal Homolog pal-1 Is Required for Posterior Patterning in Caenorhabditis elegans Embryogenesis. Dev Biol. 229(1):71–88. doi:10.1006/dbio.2000.9977.

Evans TC, Crittenden SL, Kodoyianni V, Klmble J. 1994. Translational Control of Maternal glp-1 mRNA Establishes an Asymmetry in the C. elegans Embryo. Cell. 77(2):183–94. doi:10.1016/0092-8674(94)90311-5.

Ewe CK, Sommermann EM, Kenchel J, Flowers SE, Maduro MF, Joshi PM, Rothman JH. 2022. Feedforward regulatory logic controls the specification-to-differentiation transition and terminal cell fate during Caenorhabditis elegans endoderm development. Development. 149(12):dev200337. doi:10.1242/dev.200337.

Fukushige T, Brodigan TM, Schriefer LA, Waterston RH, Krause M. 2006. Defining the transcriptional redundancy of early bodywall muscle development in C. elegans : evidence for a unified theory of animal muscle development. Genes Dev. 20(24):3395–3406. doi:10.1101/gad.1481706.

Fukushige T, Krause M. 2005. The myogenic potency of HLH-1 reveals wide-spread developmental plasticity in early C. elegans embryos. Development. 132(8):1795–1805. doi:10.1242/dev.01774.

Gilbert SF, Barresi MJF. 2016. Developmental biology. Sunderland, Massachusetts U.S.A.: Sinauer Associates, Inc.

Good K, Ciosk R, Nance J, Neves A, Hill RJ, Priess JR. 2004. The T-box transcription factors TBX-37 and TBX-38 link GLP-1/Notch signaling to mesoderm induction in C. elegans embryos. Development. 131(9):1967–1978. doi:10.1242/dev.01088.

Hartwell LH, Hopfield JJ, Leibler S, Murray AW. 1999. From molecular to modular cell biology. Nature. 402(S6761):C47–C52. doi:10.1038/35011540.

Hoege C, Hyman AA. 2013. Principles of PAR polarity in Caenorhabditis elegans embryos. Nat Rev Mol Cell Biol. 14(5):315–322. doi:10.1038/nrm3558.

Hunter CP, Kenyon C. 1996. Spatial and Temporal Controls Target pal-1 Blastomere-Specification Activity to a Single Blastomere Lineage in C. elegans Embryos. Cell. 87(2):217–226. doi:10.1016/S0092-8674(00)81340-9.

Kalb JM, Lau KK, Goszczynski B, Fukushige T, Moons D, Okkema PG, McGhee JD. 1998. pha-4 is Ce-fkh-1, a fork head /HNF-3α,β,γ homolog that functions in organogenesis of the C. elegans pharynx. Development. 125(12):2171–2180. doi:10.1242/dev.125.12.2171.

Kiefer JC, Smith PA, Mango SE. 2007. PHA-4/FoxA cooperates with TAM-1/TRIM to regulate cell fate restriction in the C. elegans foregut. Dev Biol. 303(2):611–624. doi:10.1016/j.ydbio.2006.11.042.

Kimble J, Nüsslein-Volhard C. 2022. The great small organisms of developmental genetics: Caenorhabditis elegans and Drosophila melanogaster. Dev Biol. 485:93–122. doi:10.1016/j.ydbio.2022.02.013.

Kipreos ET, Van Den Heuvel S. 2019. Developmental Control of the Cell Cycle: Insights from Caenorhabditis elegans. Genetics. 211(3):797–829. doi:10.1534/genetics.118.301643.

Labouesse M, Hartwieg E, Horvitz HR. 1996. The Caenorhabditis elegans LIN-26 protein is required to specify and/or maintain all non-neuronal ectodermal cell fates. Development. 122(9):2579–2588. doi:10.1242/dev.122.9.2579.

Liu J, Murray JI. 2023. Mechanisms of lineage specification in Caenorhabditis elegans. Seydoux G, editor. GENETICS. 225(4):iyad174. doi:10.1093/genetics/iyad174.

Ma X, Zhao Z, Xiao L, Xu W, Kou Y, Zhang Y, Wu G, Wang Y, Du Z. 2021. A 4D single-cell protein atlas of transcription factors delineates spatiotemporal patterning during embryogenesis. Nat Methods. 18(8):893–902. doi:10.1038/s41592-021-01216-1.

Maduro MF. 2009. Structure and evolution of the C. elegans embryonic endomesoderm network. Biochim Biophys Acta BBA - Gene Regul Mech. 1789(4):250–260. doi:10.1016/j.bbagrm.2008.07.013.

Maduro MF. 2010. Cell fate specification in the C. elegans embryo. Dev Dyn. 239(5):1315–1329. doi:10.1002/dvdy.22233.

Maduro MF, Hill RJ, Heid PJ, Newman-Smith ED, Zhu J, Priess JR, Rothman JH. 2005. Genetic redundancy in endoderm specification within the genus Caenorhabditis. Dev Biol. 284(2):509–522. doi:10.1016/j.ydbio.2005.05.016.

Mello CC, Draper BW, Prless JR. 1994. The maternal genes apx-1 and glp-1 and establishment of dorsal-ventral polarity in the early C. elegans embryo. Cell. 77(1):95–106. doi:10.1016/0092-8674(94)90238-0.

Mello CC, Schubert C, Draper B, Zhang W, Lobel R, Priess JR. 1996. The PIE-1 protein and germline specification in C. elegans embryos. Nature. 382(6593):710–712. doi:10.1038/382710a0.

Murray JI, Boyle TJ, Preston E, Vafeados D, Mericle B, Weisdepp P, Zhao Z, Bao Z, Boeck M, Waterston RH. 2012. Multidimensional regulation of gene expression in the C. elegans embryo. Genome Res. 22(7):1282–1294. doi:10.1101/gr.131920.111.

Owraghi M, Broitman-Maduro G, Luu T, Roberson H, Maduro MF. 2010. Roles of the Wnt effector POP-1/TCF in the C. elegans endomesoderm specification gene network. Dev Biol. 340(2):209–221. doi:10.1016/j.ydbio.2009.09.042.

Packer JS, Zhu Q, Huynh C, Sivaramakrishnan P, Preston E, Dueck H, Stefanik D, Tan K, Trapnell C, Kim J, et al. 2019. A lineage-resolved molecular atlas of C. elegans embryogenesis at single-cell resolution. Science. 365(6459):eaax1971. doi:10.1126/science.aax1971.

Phillips BT, Kidd AR, King R, Hardin J, Kimble J. 2007. Reciprocal asymmetry of SYS-1/β-catenin and POP-1/TCF controls asymmetric divisions in Caenorhabditis elegans. Proc Natl Acad Sci. 104(9):3231–3236. doi:10.1073/pnas.0611507104.

Priess JR. 2005. Notch signaling in the C. elegans embryo. WormBook.:1–16. doi:10.1895/wormbook.1.4.1.

Quake SR. 2022. A decade of molecular cell atlases. Trends Genet. 38(8):805– 810. doi:10.1016/j.tig.2022.01.004.

Regev A, Teichmann SA, Lander ES, Amit I, Benoist C, Birney E, Bodenmiller B, Campbell P, Carninci P, Clatworthy M, et al. 2017. The Human Cell Atlas. eLife. 6:e27041. doi:10.7554/eLife.27041.

Rose L, Gönczy P. 2014. Polarity establishment, asymmetric division and segregation of fate determinants in early C. elegans embryos. WormBook.:1–43. doi:10.1895/wormbook.1.30.2.

Rothman J, Jarriault S. 2019. Developmental Plasticity and Cellular Reprogramming in Caenorhabditis elegans. GENETICS. 213(3):723–757. doi:10.1534/genetics.119.302333.

Schedl T, editor. 2013. Germ Cell Development in C. elegans. New York, NY: Springer New York (Advances in Experimental Medicine and Biology). [accessed 2024 Aug 27]. https://link.springer.com/10.1007/978-1-4614-4015-4.

Seydoux G, Fire A. 1994. Soma-germline asymmetry in the distributions of embryonic RNAs in Caenorhabditis elegans. Development. 120(10):2823–2834. doi:10.1242/dev.120.10.2823.

Sulston JE, Schierenberg E, White JG, Thomson JN. 1983. The embryonic cell lineage of the nematode Caenorhabditis elegans. Dev Biol. 100(1):64–119. doi:10.1016/0012-1606(83)90201-4.

Tavernier N, Labbé JC, Pintard L. 2015. Cell cycle timing regulation during asynchronous divisions of the early C. elegans embryo. Exp Cell Res. 337(2):243–248. doi:10.1016/j.yexcr.2015.07.022.

Tintori SC, Osborne Nishimura E, Golden P, Lieb JD, Goldstein B. 2016. A Transcriptional Lineage of the Early C. elegans Embryo. Dev Cell. 38(4):430–444. doi:10.1016/j.devcel.2016.07.025.

Van Den Heuvel S. 2005. Cell-cycle regulation. WormBook.:1–16. doi:10.1895/wormbook.1.28.1.

Waddington CH. 1957. The strategy of the genes: a discussion of some aspects of theoretical biology. Routledge. [accessed 2024 Aug 27]. https://www.taylorfrancis.com/books/9781317657552.

Wang J, Yuan R, Zhu X, Ao P. 2020. Adaptive Landscape Shaped by Core Endogenous Network Coordinates Complex Early Progenitor Fate Commitments in Embryonic Pancreas. Sci Rep. 10(1):1112. doi:10.1038/s41598-020-57903-0.

Wolpert L, Tickle C, Arias AM, Lawrence P, Locke J. 2019. Principles of development. Oxford, United Kingdom: Oxford University Press.

Wood WB. 1988. The nematode caenorhabditis elegans. Cold Spring Harbor Laboratory.

Yuan R, Zhu X, Wang G, Li S, Ao P. 2017. Cancer as robust intrinsic state shaped by evolution: a key issues review. Rep Prog Phys. 80(4):042701. doi:10.1088/1361-6633/aa538e.

Zhang X, Chen Y-C, Yao M, Xiong R, Liu B, Zhu X, Ao P. 2024. Potential therapeutic targets of gastric cancer explored under endogenous network modeling of clinical data. Sci Rep. 14(1):13127. doi:10.1038/s41598-024-63812-3.

Zhu J, Hill RJ, Heid PJ, Fukuyama M, Sugimoto A, Priess JR, Rothman JH. 1997. end-1 encodes an apparent GATA factor that specifies the endoderm precursor in Caenorhabditis elegans embryos. Genes Dev. 11(21):2883–2896. doi:10.1101/gad.11.21.2883.

